# Spatial scale moderates the shape of the biodiversity-disease relationship

**DOI:** 10.1101/472902

**Authors:** Fletcher W. Halliday, Jason R. Rohr

## Abstract

Diverse host communities commonly inhibit the spread of parasites in studies at small and intermediate scales, leading some to suggest that conserving biodiversity could help control infectious diseases. However, the generality of this “dilution effect” remains controversial. First, most studies assume a linear, monotonic relationship between biodiversity and disease, though the actual shape is unknown. Second, most studies are conducted at a single spatial scale, though biotic interactions are often-scale-dependent, thus spatial scale might determine the direction of biodiversity-disease relationships. Third, most studies focus only on a small range of possible diversity levels, though the direction of biodiversity-disease relationships may change outside of this range. By analyzing 231 biodiversity-disease relationships on 77 parasite species, we provide broad evidence that biodiversity-disease relationships are generally non-linear and moderated by spatial scale; biodiversity generally inhibits disease at local scales (<100 km^2^) and amplifies disease at regional scales (>1,000,000 km^2^). These effects did not depend on any tested host, parasite, or study characteristics, though the spatial scale of a study was often related to study design and parasite type, highlighting the need for additional multiscale research. Few studies were missing substantial data at low diversity, but missing data at low diversity could result in underreporting of amplification. Experiments might be missing data at high diversity, which could result in underreporting of dilution. Despite context-dependence in biodiversity-disease relationships, most conservation is implemented at local scales where biodiversity appears to inhibit disease and thus these results suggest that local conservation actions could reduce disease risk.

**Significance statement:** It has been suggested that diverse ecological communities limit disease spread, but the generality of this pattern is contentious. Therefore, the degree to which biodiversity conservation can limit harmful epidemics remains unresolved. We address this fundamental question by analyzing 231 published relationships between biodiversity and disease. We find evidence that most biodiversity-disease relationships are nonlinear and scale-dependent with biodiversity generally associated with reduced disease at small and intermediate scales, but increased disease at large scales. Moreover, these results were generally robust to missing data at low and high biodiversity levels and variation in host, parasite, and study characteristics. This suggests that conservation efforts aimed at reducing the impacts of human and wildlife diseases will be most successful at local scales.

## Introduction

Understanding whether there is a general relationship between biodiversity and disease risk is critical for projecting and reducing the impacts of future disease outbreaks (1, 2, 11–15, 3–10). If increasing biodiversity generally reduces disease, a phenomenon coined the dilution effect, then biodiversity loss could have negative consequences for human and wildlife populations (2, 16). Concordantly, biodiversity conservation could limit the spread of unknown infectious diseases (13, 17). However, if biodiversity-disease relationships are idiosyncratic or context dependent, then biodiversity conservation could have no effect on, or, in the case of an amplification effect, even exacerbate the risk of disease to wildlife and humans (18, 19). Such context dependence in the biodiversity-disease relationship has become a major concern among disease ecologists (8, 13, 20). Consequently, the value of conserving biodiversity to protect against disease risk has been called into question (20–22).

Context dependence in the biodiversity-disease relationship can arise when the shape of the biodiversity-disease relationship is nonlinear. By definition, parasites require hosts for food and habitat. Thus, all else being equal, an increase in host biodiversity from zero hosts must initially increase the risk of disease (1, 12). However, if parasites are selected to infect the most abundant and widespread hosts or there are trade-offs between defending against parasites and host growth, reproduction, and dispersal, then communities might assemble in a manner where the first species added to communities are generally competent, disease amplifying hosts and later additions might be rarer, diluting hosts (23). If so, the initial increase in disease risk when moving from zero hosts to a few might reverse at higher diversity levels (Fig. 1a), and the skew of the biodiversity-disease relationship might affect the predominance of amplification or dilution. When biodiversity-disease relationships are left-skewed or asymptotic, amplification effects should predominate, because most increases in biodiversity will be associated with increased parasite abundance (12) (Fig. 1a). Alternatively, when biodiversity-disease relationships are right-skewed, dilution should predominate (12). Nevertheless, understanding the shape of nonlinear biodiversity-disease relationships remains a major research gap (1, 15, 20, 23).

**Fig 1.**
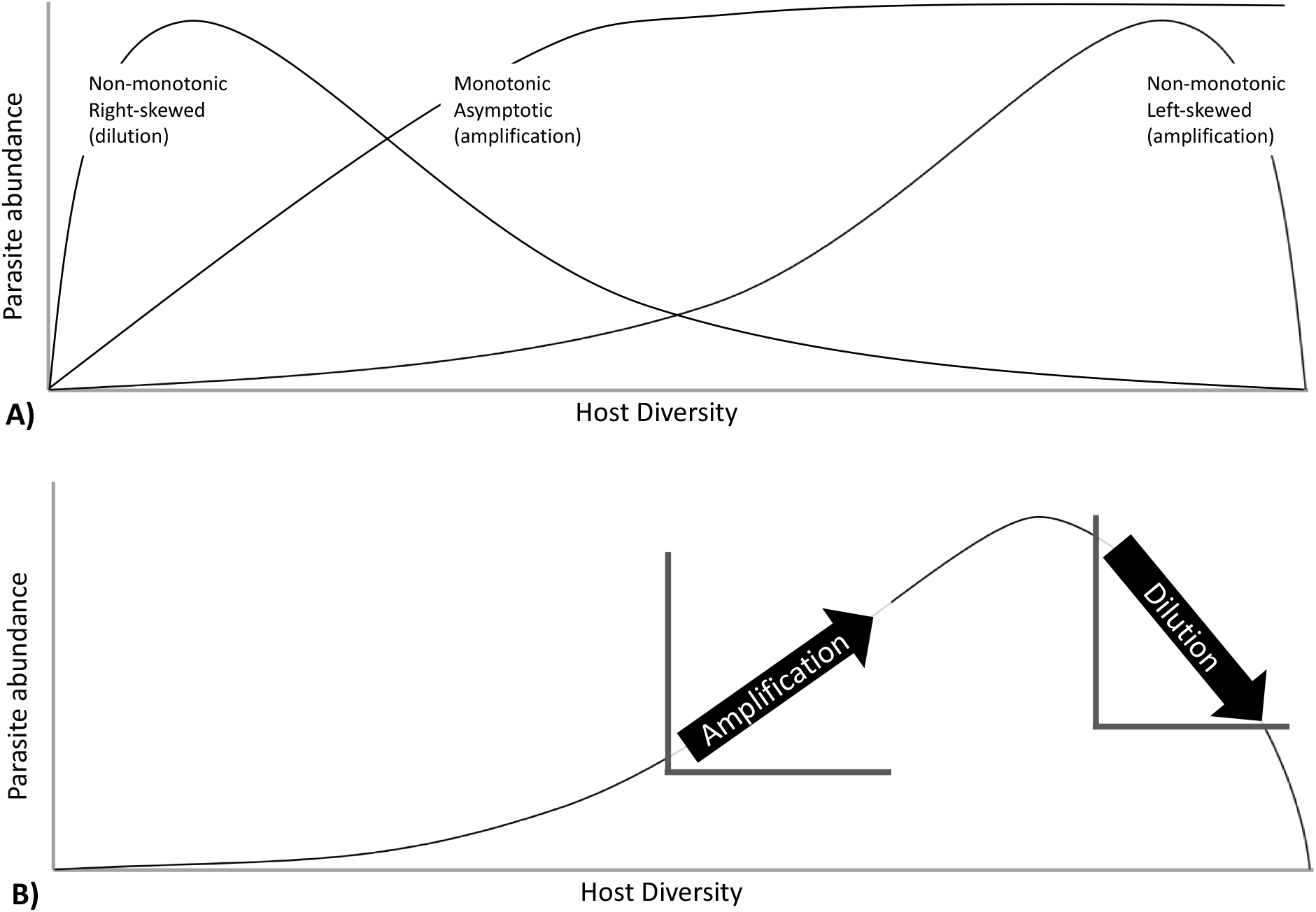
Hypothetical relationships between biodiversity and disease risk. A) A non-monotonic right-skewed distribution suggests that dilution might occur more frequently, but less intensely than amplification because the relationship is moderately negative over a greater portion of the biodiversity gradient than it is strongly positive. A non-monotonic left-skewed distribution suggest that amplification might occur more frequently but less intensely than dilution, because the relationship is moderately positive over a greater portion of the biodiversity gradient than it is strongly negative. A monotonic and asymptotic distribution suggests that amplification becomes increasingly moderate with biodiversity. B) In addition to the shape of biodiversity-disease relationships, the location on the curve where biodiversity levels are observed will also affect the likelihood and intensity of dilution and amplification. For example, in a left-skewed biodiversity-disease relationship, collecting measurements at biodiversity beyond the peak of parasite abundance could lead researchers to conclude that there is was a linear dilution effect, whereas measurements before the peak of parasite abundance would lead researchers to conclude that there was a linear amplification effect.

Where communities fall on nonlinear biodiversity-disease curves is also important. If changes in biodiversity all occur to the right or left of the peak of unimodal diversity-disease curves, then dilution or amplification, respectively, will be most common, regardless of the direction of the skew of that relationship (Fig 1b). This might create biases for both observational and manipulative studies. Observational studies might not capture low host diversity levels if they are rare in nature, which could lead researchers to spuriously conclude that there is a linear dilution effect, even though amplification might occur at non-sampled low levels of host diversity (15, 20) (Fig 1b). In contrast, manipulative studies might include mostly low levels of biodiversity because of the logistical challenges of collecting sufficient numbers of many host species for an experiment; this could potentially bias results towards amplification. The former bias can be easily detected by simply quantifying the minimum diversity level in observational studies, whereas the latter bias of experimental studies might be more difficult to assess because, in most systems, the distribution of natural levels of host diversity is unknown.

Context dependence in the biodiversity-disease relationship may also arise when the direction of the biodiversity-disease relationship depends on the spatial scale of observation (4, 7, 20). Local processes influence the abundance of species at relatively small spatial scales, while regional processes influence the distributions of species across large spatial extents (24). Relying on this well-characterized ecological phenomenon, it has been proposed that biodiversity-disease relationships should be strongest at local scales, where biotic interactions are most likely to occur, and should weaken or even reverse at larger scales, where abiotic factors like climate may cause the distributions of hosts and parasites to covary (23, 25). Moreover, whether hosts can dilute disease might be more observable at small scales where encounter reduction can occur, while the amplifying effect of hosts might only be observable at larger temporal and spatial scales (26). Even though theory indicates that spatial scale can moderate biodiversity-disease relationships, and biodiversity-disease studies have occurred from global to local scales (3, 27, 28), few studies have been conducted across multiple spatial scales. Thus the degree to which biodiversity-disease relationships are moderated by spatial scale remains largely untested (but see 25).

By quantifying the shape and direction of 231 published biodiversity-disease relationships, this study aimed at testing three contingencies in the biodiversity-disease relationship. Specifically, we tested whether biodiversity-disease relationships are generally (a) non-linear, (b) moderated by spatial scale, and (c) sensitive to missing data at low and high diversity. Our results indicate that biodiversity-disease relationships are generally non-linear, that dilution most commonly occurs at small (i.e., local) scales and amplification most commonly occurs at large (i.e., regional) scales, that few studies are missing substantial data at low diversity, but that missing data at low diversity could potentially result in the underreporting of amplification effects, and that experimental studies might be missing data at high diversity levels that could potentially result in underreporting of dilution effects.

## Results and Discussion

### What is the shape of the biodiversity-disease relationship?

First, we tested whether the published relationship between biodiversity and disease was linear or nonlinear by comparing intercept-only, linear, second-order, and third-order polynomial regression models for all biodiversity-disease relationships, selecting the best-fitting model using AIC (Table S1). Importantly, these models were only based on the data presented in each study and thus did not constrain the biodiversity-disease relationship to the origin (see “Do *missing data at low and high diversity bias studies to report dilution effects*?” section below). Out of the 231 studies that included more than three levels of biodiversity, 63% were best fit by a linear, second-order, or third-order polynomial model (i.e., exhibited a relationship between biodiversity and disease). Of these studies, biodiversity-disease relationships were most commonly non-linear, as predicted. More specifically, 60% exhibited non-linear relationships (either second or third-order polynomial), while 8% exhibited linear amplification effects, and 32% exhibited linear dilution effects.

While comparing regression models identified many non-linear biodiversity-disease relationships, this approach is constrained by the functional form of each regression model. In other words, we are only able to detect non-linear relationships where those relationships were best fit by second- or third-order polynomials. To relax this constraint, we used Spearman rank correlation tests (not constrained to pass through the origin), which make no assumption about the underlying distribution of the data nor the linearity of the relationship between variables, and are therefore not constrained by the functional form of the biodiversity-disease relationship. We quantified whether each biodiversity-disease relationship was monotonic and positive (disease increases, but may level off, as diversity increases), monotonic and negative (disease decreases but may level off as diversity increases), or non-monotonic (disease increases with diversity at low levels, but eventually decreases at high enough diversity; Fig. 1a). The estimated Spearman rank correlation coefficient (ρ) approaches one for monotonic, positive relationships, and approaches negative one for monotonic, negative relationships. We therefore used p to define monotonic amplification (ρ>0, *p*<0.05), monotonic dilution (ρ<0, *p*<0.05), and non-significant or non-monotonic relationships (*p*>0.05). Consistent with the previous analysis, 15% of the 231 relationships exhibited monotonic amplification effects, 31% exhibited monotonic dilution effects, and 54% exhibited non-significant or non-monotonic relationships.

Given that non-linear and non-monotonic biodiversity-disease relationships are most common and that amplification effects might predominate when these relationships are left-skewed or asymptotic, whereas dilution might predominate when they are right-skewed (12), we next assessed the skew of each biodiversity-disease relationship. To do so, we fit a smoothing spline to each published biodiversity-disease relationship, that was not constrained to pass through the origin, and then calculated Pearson’s skewness from the shape of the estimated curve, excluding studies where there was no relationship (i.e., where the slope of the curve was not significantly different from zero; *n*=29). As expected, Pearson’s skewness and Spearman rank correlation were in agreement when studies exhibited monotonic biodiversity-disease relationships. Specifically, studies exhibiting monotonic dilution effects were significantly right-skewed (*p*<0.001), and studies exhibiting monotonic amplification effects were significantly left-skewed (*p*<0.001; Fig. 2). Studies exhibiting non-significant or non-monotonic relationships based on Spearman rank correlation were not significantly skewed (*p*=0.59), indicating that non-monotonic biodiversity-disease relationships, on average, showed similar levels of amplification or dilution. These results were qualitatively similar for the analysis comparing intercept-only, linear, second-order, and third-order polynomial regression models (Fig. S1, Fig. S2).

**Fig 2.**
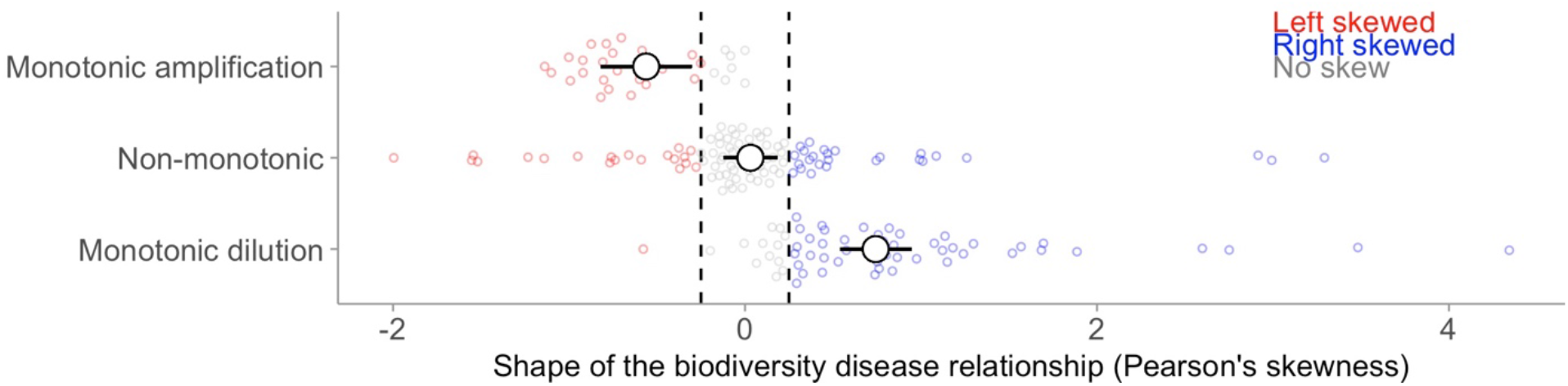
Results of the analysis comparing Spearman rank correlation to Pearson’s skewness. Points are model-estimated means and error bars are 95% confidence intervals. The colored points show the distribution of the raw data. Left-skewed relationships (Pearson’s skewness<0.25) are shown in red, right-skewed relationships (Pearson’s skewness>0.25) are shown in blue, and non-skewed relationships are shown in grey. Spearman rank correlation was strongly associated with Pearson’s skewness: monotonic amplification effects (ρ>0, p<0.05) tended to be left-skewed, monotonic dilution effects (ρ<0, p<0.05) were right skewed, and nonmonotonic relationships were not significantly skewed.

These results support the hypothesis that the shape of biodiversity-disease relationships might be nonlinear (12, 20) and might therefore have implications for biodiversity conservation. Specifically, because biodiversity-disease relationships are often nonlinear, where an individual system falls along a biodiversity gradient might influence whether that system experiences amplification or dilution. Thus, understanding how an individual conservation action will alter biodiversity can have a large impact on whether that action is expected to increase or decrease wildlife and human disease risk.

### Is the biodiversity-disease relationship moderated by spatial scale?

Second, we tested whether the shape and direction of the biodiversity-disease relationship was moderated by spatial scale, measured as the spatial extent of each study. Spatial extent represents the total area over which a study is conducted, including all measures of biodiversity and disease for a given study. Spearman’s ρ, which measures the monotonicity and direction of association between biodiversity and disease, was positively associated with spatial extent (*p*<0.001; marginal *R*^2^=0.34; Fig 3a), with monotonic dilution effects most commonly occurring at small to intermediate spatial scales and monotonic amplification effects most commonly occurring at the largest spatial scales. Incorporating the shape of non-monotonic relationships did not alter this result; Pearson’s skewness was significantly associated with spatial extent (*p*<0.001; marginal *R*^2^=0.22; Fig 3b), with right-skewed relationships (indicating more dilution) occurring at small to intermediate spatial scales and left-skewed relationships (indicating more amplification) occurring at large spatial scales.

**Fig 3.**
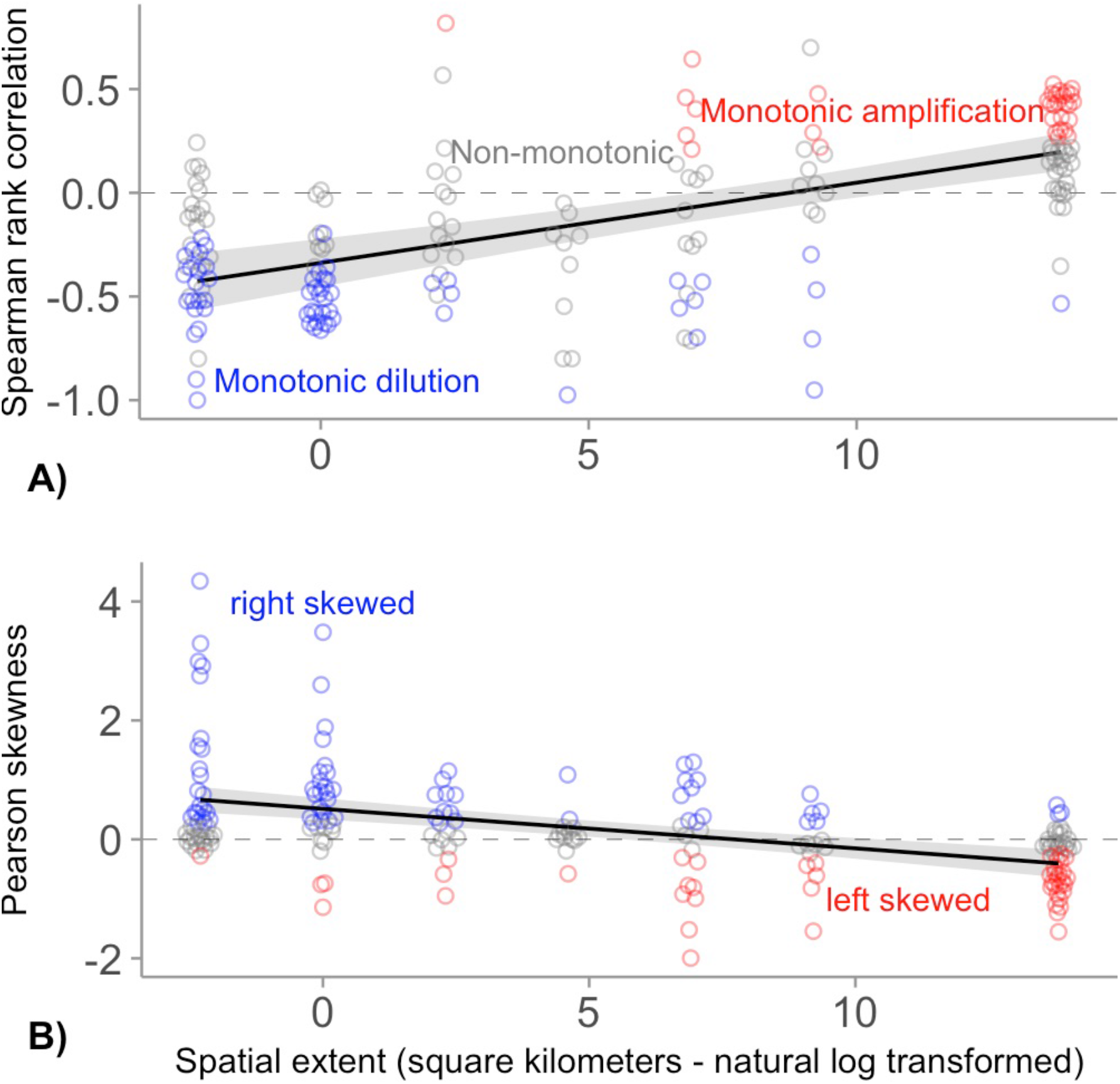
Results of the analyses relating spatial scale to the shape of the biodiversity-disease relationship. Points represent each published biodiversity-disease relationship, colored by their estimated shape (red = Monotonic amplification in panel A and left-skewed in panel B; blue = Monotonic dilution in panel A and right-skewed in panel B; grey = non-significant or nonmonotonic in panel A non-skewed in panel B). Solid lines indicate the estimated fit of a multilevel random effects model, and grey ribbons indicate the 95% confidence intervals. Spatial scale moderates the relationship between biodiversity and disease: A) Spearman rank correlation between biodiversity and disease was positively associated with spatial extent, and B) Pearson’s skewness was negatively associated with spatial extent.

Dilution generally occurred *within* an ecosystem, at spatial extents <100 km^2^ (roughly the size of a small city), whereas amplification generally occurred *across* ecosystems, in studies occupying >1,000,000 km^2^ (roughly the size of France and Spain combined). These results therefore indicate that the overall disease burden in one ecosystem can be higher than another because its native biodiversity is higher, but if this ecosystem has its biodiversity lowered, disease could still worsen. Consequently, these results indicate that within individual countries, conserving biodiversity might improve human, wildlife, and ecosystem health.

This dependence of biodiversity-disease relationships on spatial scale may be an indicator of a more general mechanism of disease amplification. Notably, comparison of biodiversity-disease relationships *within* an ecosystem often include many of the same host species (e.g., 29), whereas comparisons of biodiversity *across* ecosystems tend to include distinct sets of host species (e.g., 30). Thus, measuring the degree to which host-species turnover (β diversity) drives disease amplification could help clarify why amplification occurs at large spatial extents, and possibly help predict when amplification will be more common, in general.

These results reveal a strong, widespread association between the shape of biodiversity-disease relationships and the spatial scale of observations, supporting the hypothesis that biodiversity-disease relationships are scale-dependent (7, 23). Importantly, however, not every small-scale study exhibited dilution, nor did every large-scale study exhibit amplification. As an example, in a global survey of human disease burden, disease generally increased with increasing diversity, but human schistosomiasis was negatively correlated with diversity (30) and, at small spatial scales, biodiversity can amplify disease via a sampling effect, if species are added randomly with respect to host competence and transmission is frequency-dependent (31).

We tested whether several ecological factors could explain variation in the effect of spatial scale on the shape of biodiversity-disease relationships. Specifically, we tested whether the effect of spatial scale on biodiversity-disease relationships differed between (i) parasites that infect humans vs. wildlife, (ii) macro-vs. microparasites, (iii) parasites with complex vs. direct lifecycles, and (iv) observational vs. manipulative studies. We found no evidence that the effect of spatial scale on biodiversity-disease relationships depended on any of these factors (Table 1). Thus, the effect of spatial scale on biodiversity-disease relationships was generally robust across all ecological contexts examined. However, we encourage caution in interpreting these results, as there was multicollinearity in these analyses. Specifically, observational studies and studies of human pathogens both tended to occur at larger spatial scales than manipulative studies and studies of wildlife pathogens (Fig. S3). Consequently, we cannot rule out the possibility that these results could change if future studies filled these research gaps, allowing tests of these context dependencies to be less collinear.

**Table 1.** Models of ecological factors moderating the effect of spatial scale on biodiversity-disease relationships

|  | A) Spearman correlation coefficient |  |  | B) Pearson's skewness |  |  |
| --- | --- | --- | --- | --- | --- | --- |
|  | **DF** | **F-value** | **p-value** | **DF** | **F-value** | **p-value** |
| Spatial extent | 41.3 | 2.862 | 0.098 | 4.6 | 0.395 | 0.56 |
| Human | 65.0 | 2.171 | 0.145 | 57.6 | 0.061 | 0.81 |
| Route | 61.2 | 4.124 | 0.047 | 55.3 | 0.670 | 0.42 |
| Macroparasite | 30.5 | 1.335 | 0.257 | 12.3 | 0.152 | 0.70 |
| Manipulative | 61.2 | 0.664 | 0.418 | 14.2 | 0.020 | 0.89 |
| Spatial extent × Human | 42.5 | 2.140 | 0.151 | 15.6 | 0.007 | 0.93 |
| Spatial extent × Route | 69.9 | 2.464 | 0.121 | 87.7 | 0.523 | 0.47 |
| Spatial extent × Macroparasite | 34.3 | 0.385 | 0.539 | 23.2 | 1.041 | 0.32 |
| Spatial extent × Manipulative | 43.6 | 0.046 | 0.831 | 4.0 | 0.010 | 0.93 |

Type III Analysis of Variance Table with Satterthwaite's method
DF: Denominator degrees of freedom

### Do missing data at low and high diversity bias biodiversity-disease studies?

Finally, we tested the hypothesis that missing data at the highest and lowest diversity levels in experimental and observational studies might bias studies to more commonly report amplification and dilution effects, respectively. Experimental studies had a lower mean maximum diversity level than observational studies (experimental mean ± sd: 25 ± 10, observational mean ± sd: 76 ± 89). Thus, it appears that experimental studies are missing data at the highest diversity levels, which could bias experimental studies towards amplification effects. This result could emerge from two key differences between experiments and observational studies. First, experimentally manipulating many species is logistically challenging at high richness, potentially biasing experimental studies to include fewer total species than observational studies of equivalent size. Second, the number of species in an area is highly sensitive to the area surveyed (32), and observational studies were, on average, five orders of magnitude larger than manipulative experiments (Fig S3). Focusing on studies of comparable extent (1-10 km^2^) eliminated the difference in mean maximum diversity between experiments (29 ± 10) and observational studies (22.0 ± 10), supporting this second mechanism.

We also examined the lowest diversity levels to assess whether there was missing data at low diversity. Experimental studies had lower mean minimum diversity than observational studies (experimental mean ± sd: 1.2 ± 0.6), observational mean ± sd: 4.3 ± 7.3), which could bias observational studies towards dilution effects. However, out of 231 studies, 84% of studies included an effective species richness of two or lower. Consequently, most studies (*n*=193) were not missing substantial data at low host diversity. This result indicates that the potential for missing data at low diversity to bias the estimated relationship between biodiversity and disease is quite low.

Even though most studies were not missing substantial data at low host diversity, we still performed an additional test of the hypothesis that missing data might bias studies to more commonly report dilution effects. Here, we again quantified the skew of each biodiversity-disease relationship, this time constraining each curve to pass through the origin because if there are no hosts there cannot be any parasites. Constraining each curve to pass through the origin should reduce the estimated skew in all studies, particularly studies that found monotonic dilution effects. As predicted, constraining the curves to the origin significantly changed the shape of the average biodiversity-disease relationship (*p* = 0.001), reducing the estimated frequency of dilution effects and increasing the estimated frequency of amplification effects (Fig. 4). This result indicates that ignoring missing data may bias some studies to underreport amplification effects. However, even though constraining curves to fit through the origin shifted the estimated skew, on average, the constrained curves were only marginally significantly left-skewed (*p*=0.051; Fig. S4). Furthermore, spatial scale still significantly moderated the sign of the constrained curves, with dilution more common at small scales and amplification more common at large scales (p<0.0001; Fig. S5), and this effect was still robust to ecological characteristics of individual study systems (Table S2).

**Fig 4.**
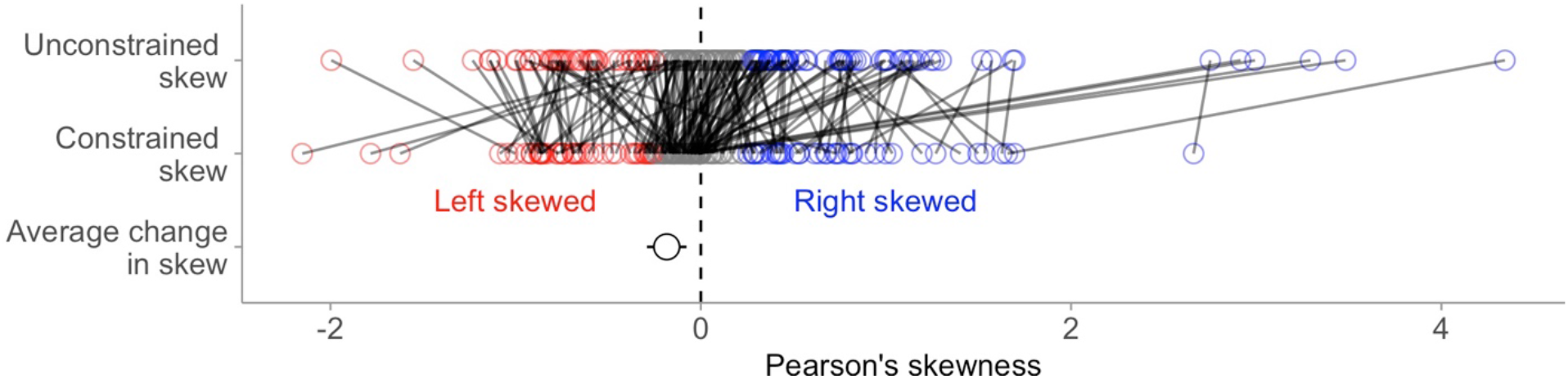
Results of constraining biodiversity-disease relationships to pass through the origin. The top two rows show Pearson’s skewness for unconstrained curves, and curves that were constrained to pass through the origin, with each study connected by a solid line. Left-skewed relationships (Pearson’s skewness<0.25) are shown in red, right-skewed relationships (Pearson’s skewness>0.25) are shown in blue, and non-skewed relationships are shown in grey. The bottom row shows the model-estimated effect of constraining the curves to pass through the origin, with the point indicating the model-estimated mean, and error bars showing the 95% confidence interval. On average, constraining curves to pass through the origin results in a more left-skewed relationship between biodiversity and disease.

These results indicate that scale-dependence of the biodiversity-disease relationship is robust to missing data at low diversity levels. The robustness of this scale-dependence may be a product of the underlying shape of biodiversity-disease relationships. Studies that found monotonic amplification effects were unlikely to be altered by missing data at low diversity (Fig 1a). Conversely, studies that found monotonic dilution had higher potential to be altered by missing data at low diversity. However, 66 of the 72 studies showing monotonic dilution effects included an effective species richness of two or lower. Thus, regardless of the shape of the relationship between the origin and the point of peak parasite abundance, the area in which amplification could occur was generally quite small. We therefore conclude that although the biodiversity-disease relationship can take on many forms, and its form may depend on a nonlinearity that is driven by parasite extinction at low host diversity, such nonlinearities are unlikely to alter a general and common phenomenon: dilution effects most commonly occur at local scales and amplification effects most commonly occur at regional scales.

Together, these results indicate that the scale-dependence of biodiversity-disease relationships might be robust to missing data at low diversity levels and to ecological characteristics of individual studies. However, we are unable to test whether missing data at high diversity might bias experimental studies to more commonly observe amplification. This bias of experimental studies is difficult to assess because, in most systems, the distribution of natural levels of host diversity is unknown. Furthermore, a major limitation to this analysis is the lack of empirical studies conducted across spatial scales and ecological conditions using the same methodologies. We are hopeful that as large-scale replicated studies, such as the Nutrient Network (33) and National Environmental Observatory Network (34), become more widespread, the quality of data at the largest spatial scales will improve. These results also highlight the need to consider whether the scales of conservation actions and public health interventions are appropriate to influence biodiversity-disease relationships in a way that will benefit humans (3, 27, 30). In general, these results suggest that biodiversity conservation can be beneficial to human health when conducted at small or intermediate scales, which are the scales at which they are most commonly implemented (35, 36).

Understanding how biodiversity alters infectious diseases remains a critical frontier in disease ecology (17, 23, 37) as human activities continue to alter global biodiversity (38, 39), and disease outbreaks continue to increase (40–42). This study provides quantitative evidence that the relationship between biodiversity and disease is non-linear and scale-dependent. This general pattern indicates that biodiversity loss could exacerbate disease outbreaks at the scales in which humans are most likely to encounter disease, and highlights important scales in which biodiversity conservation might be most useful for minimizing and mitigating these consequences.

## Materials and Methods

### Data compilation

This study aimed to analyze the shape of every published relationship between host diversity and the abundance of parasites. We updated the list of studies from Civitello et al. (8) to include studies published between 2014 and 2018, by repeating their original search criteria. Specifically, we searched the Web of Science for several combinations of search terms: parasite, pathogen, diversity, richness, evenness, dilution effect, and amplification effect (the final search was conducted in June 2018). We identified additional papers by searching the literature cited sections of these articles and by searching Web of Science for all papers citing Civitello et al (8), including those critical of the dilution effect hypothesis. We included observational and experimental studies in lab and field environments.

### Selection criteria and data collection

We only included studies that measured parasite abundance or prevalence at more than two host diversity levels. We included studies that reported infection prevalence, mean parasite load, density of infected vectors, or percent diseased tissue, because these quantities are the most relevant metrics of disease risk for microparasites, macroparasites, vector-borne parasites, and plant parasites, respectively. We did not standardize parasite abundance, and therefore did not compare parasite abundance among studies. Host biodiversity was reported as species richness, Simpson’s diversity index (J), or Shannon’s diversity index (H). We standardize across these measures to facilitate comparisons across studies by transforming diversity into the effective number of species, following Jost (43). In experiments, estimated diversity included all taxa added by the experimenters, while the diversity estimate in observational studies was limited to a focal taxonomic or functional group of host species, defined in the primary study (e.g., herbaceous plants, trees, birds, or small mammals).

We extracted data from text and tables manually and from figures using WebPlotDigitizer version 4.1 (44), and recorded other data relating to the biology or methodology of each study. For all studies, we recorded parasite and host taxa, type of parasite (infecting only wildlife or also infecting humans), focal host species, associated species (i.e., additional species whose presence may dilute or amplify parasite abundance, operationally defined as “potential diluters”), the diversity (e.g., richness) in the treatments (or in the field survey), parasite functional group (macroparasite vs. microparasite), parasite lifecycle (complex vs. direct), and study design (manipulative vs. observational). Spatial extent was quantified as the area (expressed in square kilometers) over which all biodiversity estimates were compared in a given study. Studies rarely provided an exact value for spatial extent. Because a value for spatial extent was rarely provided, and spatial extents varied by six orders of magnitude, we estimated the extent of each survey to the nearest order of magnitude rather than attempting to assign a specific spatial extent for each study. For example, we assigned studies a value of 0.1 if the extent was less than 1 km^2^, and a value of 1 if the extent was greater than 1 km^2^, but less than 10 km^2^, etc.

### Assessing the shape of the biodiversity-disease relationship

We first quantified whether each biodiversity-disease relationship was linear or nonlinear by comparing a series of regression models using the lm and AIC functions in R version 3.5.0 (45) (see main text for methods). Four studies included fewer than five host diversity levels and were therefore not tested using a third-order polynomial. Next, we quantified the monotonicity and direction of each biodiversity-disease relationship using Spearman rank correlations (see main text for methods). We then assessed the skew of each biodiversity-disease relationship using R package cobs (46) to fit an unconstrained spline to the biodiversity-disease relationship, limited to a maximum of four knots to prevent overfitting. This approach to fitting an unconstrained curve makes no assumptions about the underlying shape of the relationship between biodiversity and disease. We transformed the predicted curve into a frequency distribution, assigning any negative value (occurring in 19 regressions) to zero, and then calculated Pearson’s skewness. A right-skewed relationship (Pearson skewness > 0.25) indicates that most of the data falls in the area where dilution is observed, while a left-skewed relationship (Pearson skewness < −0.25) indicates the possibility for measured or unmeasured amplification effects. To assess whether missing data at low diversity could bias the estimated shape of the biodiversity-disease relationship, we constrained curves to pass through the origin and again calculated the skewness of each curve. Specifically, to fit qualitatively constrained quantile (CQ) smoothing splines (47), we added a value at the origin for each data set, corresponding to a situation in which there is no host diversity, generated a constraint matrix to force the line through the origin, and then fit the curve, limiting the maximum number of knots in the curve to three to prevent overfitting.

We omitted studies with fewer than four unique measures of host diversity for Spearman rank correlations and unconstrained splines and fewer than three unique measures of host diversity for CQ splines. Twenty-nine of the unconstrained splines (*n*=231) and 39 of the CQ splines (*n*=243) showed no relationship between biodiversity and disease (e.g., a fit with a slope of zero), resulting in no estimate of Pearson’s skewness. This resulted in 231 estimates of ρ, 202 estimates of skew from unconstrained splines, and 204 estimates of skew from CQ splines.

### Data analysis

All analyses were carried out in R version 3.5.0 (45). We constructed multilevel random effects models using the lmer function in R packages lme4 (48) and lmerTest (49). We accounted for nonindependence arising from multiple measures from the same observational units in the same year by including such non-independent surveys as random intercepts in each model.

Using the model described above, we first tested whether studies exhibiting no relationship, linear amplification, linear dilution, a unimodal relationship or a third-order polynomial relationship predicted Spearman rank correlation and Pearson’s skewness. We next verified that studies exhibiting monotonic dilution, monotonic amplification, and non-monotonic relationships (categorized using the Spearman rank correlation) predicted Pearson’s skewness. Next, we tested whether the Spearman rank correlation coefficient between biodiversity and disease or Pearson’s skewness were influenced by spatial extent by fitting two separate models, each with one response (ρ or skew) and one predictor (extent).

We then tested for context dependence in the spatial moderation of dilution effects. To test for context dependence, we fit the same two models, but included a two-way interaction between spatial extent and four binary factors that might explain variation in the effects of scale on the biodiversity-disease relationship: parasite functional group (macroparasite vs. microparasite), parasite lifecycle (complex vs. direct), study design (manipulative vs. observational), and parasite type (infects humans vs. infects only wildlife).

We next tested whether missing data at low and high diversity might bias studies to more commonly report amplification and dilution effects. We quantified the maximum and minimum diversity level of each study and compared whether the mean maximum and mean minimum diversity level differed between experiments and observational studies. Because the species-area relationship is nonlinear and sample area was highly variable across studies, we compared minimum and maximum diversity across studies qualitatively rather than quantitatively.

To quantitatively test whether missing data at low host diversity could bias studies to more commonly report dilution effects, we tested whether constraining the curves to pass through the origin altered the predicted skew. Specifically, we calculated the difference in skew between constrained and unconstrained curves and then performed an intercept-only model on this value, where an estimate significantly lower than zero would indicate that constraining the curve favored amplification, and an estimate significantly higher than zero would indicate that constraining the curve favored dilution. Finally, we analyzed whether spatial scale moderated the shape of the biodiversity-disease relationship when curves were constrained to pass through the origin. Here, we fit a model of Pearson’s skewness and spatial extent and then performed the same test of context dependence on the model that was performed before.

## Acknowledgements

We are thankful to D. Civitello, R.W. Heckman, G. Legault, C.E. Mitchell, J. Umbanhowar, and members of the Rohr lab for helpful discussions on data analysis and interpretation. C.E. Mitchell, R. Poulin, H. Young, and S. Zhou provided raw data from published manuscripts. This research was supported by grants from the National Science Foundation (EF-1241889), National Institutes of Health (R01GM109499, R01TW010286), US Department of Agriculture (NRI 2006-01370, 2009-35102-0543), and US Environmental Protection Agency (CAREER 83518801) to J. R. Rohr.

